# A local scale evaluation of spatial sampling bias in the Atlas of Australian Birds

**DOI:** 10.1101/2021.08.25.457648

**Authors:** Stephen L. Totterman

## Abstract

The reliability of ‘citizen science’ datasets where volunteers are free to choose sampling locations is not clear. This study examined local (‘patch’) scale spatial sampling patterns in the Atlas of Australian Birds and then compared reporting rates, *i.e*. the proportion of sampling units in which a given species was present, from a sample of atlas points with those from a regular sample. Three sites that have been were surveyed sequentially between January–May 2017: Killawarra Forest, Victoria, Coolah Tops National Park and Pilliga Nature Reserve, New South Wales. Spatial bias in atlas sampling patterns was evident as clusters at tourist areas and special habitat features and linear patterns along roads and creek lines. Atlas samples overestimated reporting rates for species with spatial distributions that were concordant with those sampling patterns and *vice versa*. At least two-fold differences in atlas/regular sample reporting rate ratios were detected for between 13–15% of non-rare species (with reporting rates ≥ 0.08). Concerns are raised that spatial sampling bias is common in the atlas and affects a variety of species, that popular sites may not be representative of habitat patches and that a large proportion of surveys are being filtered out in data analyses.

## INTRODUCTION

The analysis of large, volunteer-collected, datasets is a growing field in conservation science (*e.g*. Cunningham and Olsen 2009; Szabo *et al*. 2010; Szabo *et al*. 2012). However, the reliability of unstructured surveys is uncertain (Lemeshow and Levy 1999; Anderson 2001).

Spatial and temporal sampling biases have been recognised in the Atlas of Australian Birds (hereafter ‘the atlas’) (Barrett *et al*. 2003; BirdLife Australia 2015). Two primary causes of sampling bias are ‘convenience sampling’ and ‘subjective’ or ‘targeted sampling’ (Anderson 2001). Convenience sampling is the selection of more accessible locations and/or times, *e.g*. along roadways. Targeted sampling is the selection of locations and/or times where abundance and/or species richness is known to be high, *e.g*. at dams. Sampling bias can result in overestimation of abundance for species with distributions that are concordant with sampling patterns and *vice versa*.

A previous, regional scale evaluation compared atlas reporting rates, *i.e*. the proportion of sampling units in which a given species was present, with those from a structured survey, in the Mount Lofty Ranges, South Australia (hereafter ‘the Mount Lofty Ranges evaluation’) (Szabo *et al*. 2012). Bivariate regression results showed strong agreement in reporting rates (intercept ∼ 0 and slope ∼ 1), however those results are rather meaningless because the same 20-minute, two-hectare search method was used in both surveys and so systematic bias (method bias) was small relative to bias for individual species, *i.e*. differences in spatial and temporal sampling effort and observer effects that can result in variable bias for individual species.

Individual species results from the Mount Lofty Ranges evaluation showed at least two-fold differences for 17 of the 61 species (28%) but those were disregarded as mostly species using edges and open habitats (Szabo *et al*. 2012). Actually, there were at least two-fold differences for seven of 27 species (26%) that were classified as dependent on native vegetation, which is practically equal to 10 of 34 species (29%) that were classified as not dependent on native vegetation. Also, the list of species not dependent on native vegetation included the Crimson Rosella *Platycercus elegans* and Superb Fairy-wren *Malurus cyaneus*, which were two of the most common birds in woodlands (Szabo *et al*. 2011), as well as several other indicator species for dry sclerophyll woodland/forest (BirdLife Australia 2015). Large sample sizes were a strength of the Mount Lofty Ranges evaluation, with 554 atlas surveys and 3877 structured surveys, and large bias results for individual species cannot be dismissed as sampling variability.

This study performed a local scale evaluation of spatial sampling bias in the atlas in three eucalypt forests. Atlas data were private and were not requested. Instead, atlas survey points were resurveyed simultaneous with a regular sample of points. This approach avoided temporal and observer effects and focused comparisons on spatial sampling bias.

## METHODS

### Study sites

Suitable study sites were contiguous patches of open, eucalypt forest with at least 100 preexisting survey points in the atlas to sample from.

Killawarra Forest (36°13’S, 146°11’E) is a box-ironbark forest remnant in the Ovens-Murray region of Victoria, 18 km north-west of Wangaratta (Fig. 1a). The site area is 3209 Ha and it is reasonably flat, 150–250 m asl. There are several intermittent creeks across the site, that were dry during the survey, and several dams. Open box, stringybark and ironbark eucalypt forest with a ground layer of herbs and grasses grows on the ridge lines. Box woodland with an open mid-storey of wattles and a ground layer of herbs and grasses grows on the flats and along creek lines. The site was crisscrossed with many roads and tracks, most of which were rarely trafficked. There was a camp ground in the middle of the forest that was saw few visitors. Killawarra was surveyed between 28 February and 18 March 2017.

**Figure 1.**
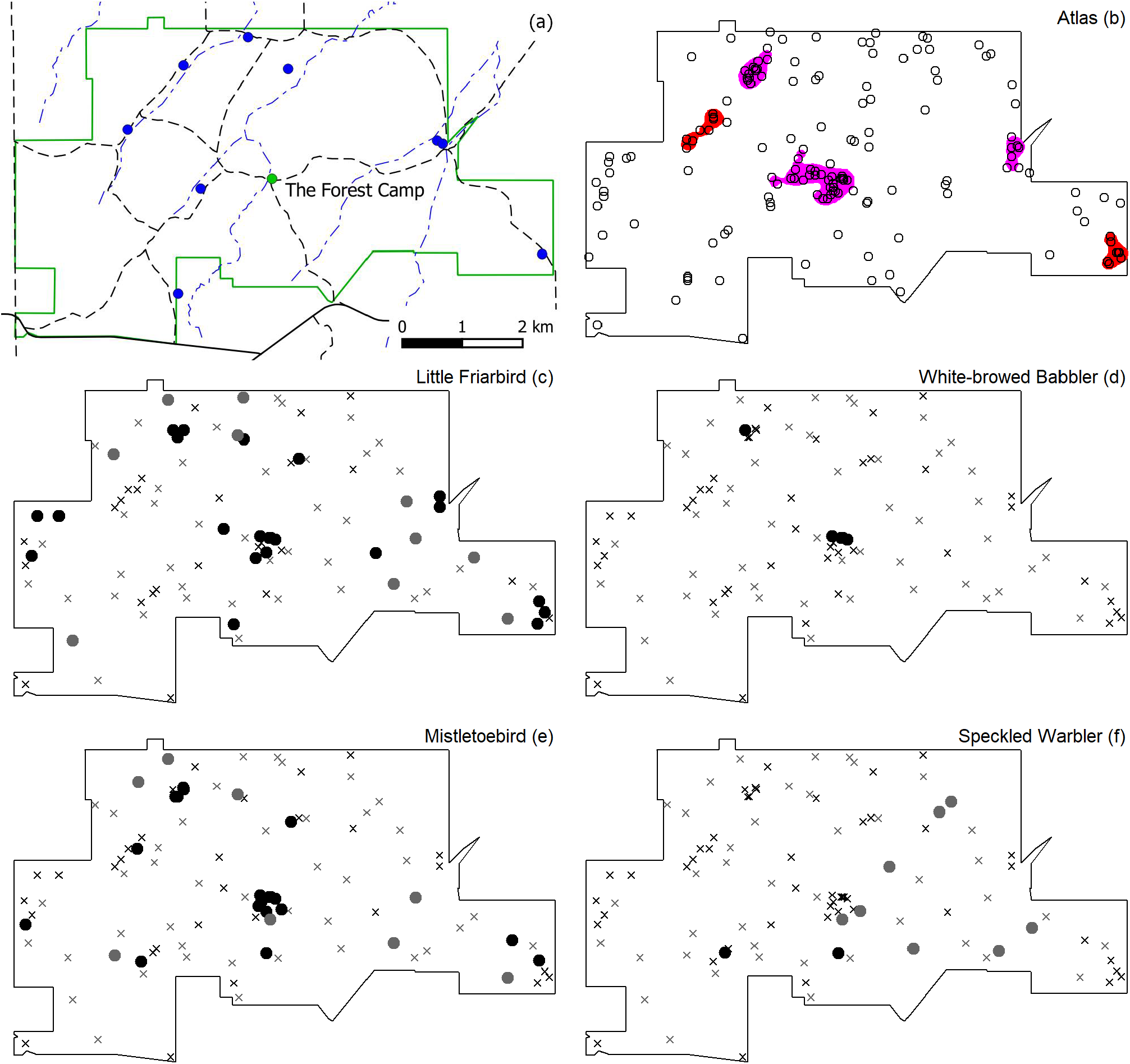
Killawarra map (a), atlas spatial sampling pattern (b) (*n* = 154) and species distribution examples (c–f). Solid and dashed black lines in (a) are sealed and unsealed roads respectively (minor tracks are not shown), blue lines are intermittent creek lines and blue circles are dams. Coloured areas in (b) are clusters that were defined using a kernel density and 50% isopleth. Magenta areas correspond to shared sites in the atlas (BirdLife Australia 2021). Black and grey circles in (c–f) are atlas sample and regular sample records respectively and crosses are species absences (total *n* = 50 in each sample).

Coolah Tops National Park (31°44’S, 150°01’E) is located on an elevated plateau, 1000–1200 mASL, at the western end of the Liverpool Range, 32 km north-east of Coolah, NSW (Fig. 2a). The park area is 14097 Ha however sampling was restricted to 10113 Ha of plateau, mostly above 1000 m asl, where most of the preexisting atlas survey points were and to avoid the rugged flanks of the range and drier forest and woodlands of the surrounding Central West Slopes and Plains region. Open stringybark and gum eucalypt forests grow on the plateau, often with a grassy ground layer. There are numerous streams draining the plateau. A dense shrub mid-storey was common along creeks and along the margins of grassy swamps. Most of the roads and tourist facilities are in the west of the forest. The Barracks and Cox’s Creek were the two most popular camp grounds. Coolah Tops was surveyed between 20 March and 11 April 2017.

**Figure 2.**
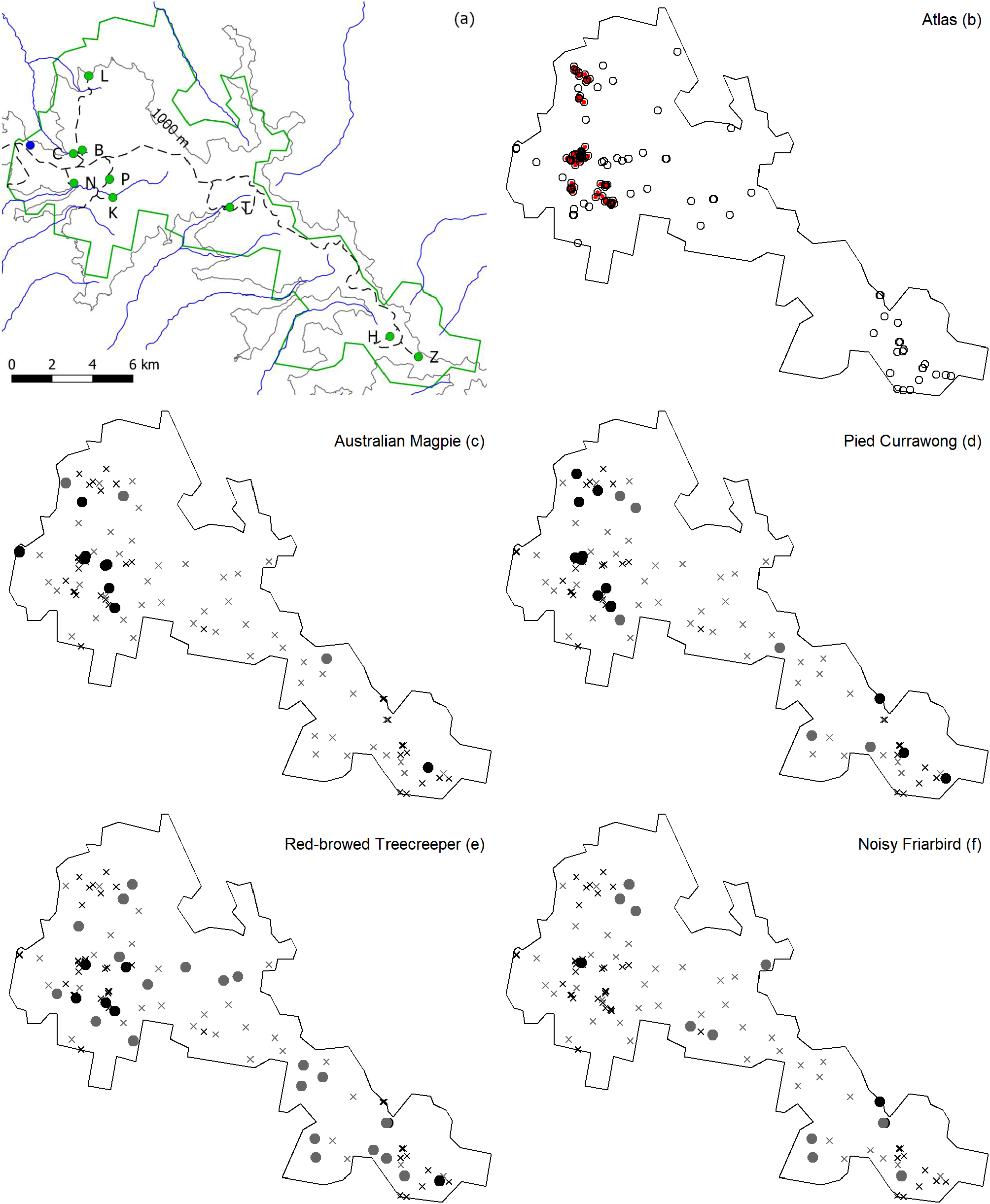
Coolah Tops map (a), atlas spatial sampling pattern (b) (*n* = 116) and species distribution examples (c–f). The green line in (a) is the Coolah Tops National Park boundary, the grey line is a 1000 m contour, dashed black lines are unsealed roads (minor tracks and management trails are not shown), blue lines are creeks and the blue circle is a dam. Green circles are tourist areas: B = The Barracks Camp, C = Cox’s Creek Camp, P = The Pines Camp, H = Cattle Creek Hut, K = Brackens Hut, L = Bundella Lookout, Z = Breeza Lookout, N = Norfolk Falls and T = Talbragar River Falls. Coloured areas in (b) are clusters that were defined using a kernel density and 75% isopleth. There were no shared sites for Coolah Tops in the atlas (BirdLife Australia 2021).

Pilliga Nature Reserve (30°53’S, 149°28’E) is at the southern end of the Pilliga Forests in the North West Slopes and Plains region of NSW, 33 km north of Coonabarabran. The study site was defined by drawing a polygon around preexisting atlas survey points (Fig. 3a). The site area is 27098 Ha and it is essentially flat, 300–500 m asl. Ironbark eucalypt and cypress pine grow on the ridge lines. Gum eucalypts grow along creek lines. The mid-storey was often quite dense with wattles and other shrubs. Mallee shrubland was found in the east of the site. The south-western corner of the site had been burnt at some unknown time preceding the survey and showed abundant regrowth. There are several intermittent creeks across the site that were dry during the survey. Borah Creek was exceptional in featuring waterholes and grassy eucalypt woodland along its banks. No dams were found within the site. The Sandstone Caves, on the southern boundary, and Yaminba Rest Area, on the Newell Highway, were the only sites frequented by visitors. Pilliga was surveyed between 15 April and 7 May 2017.

**Figure 3.**
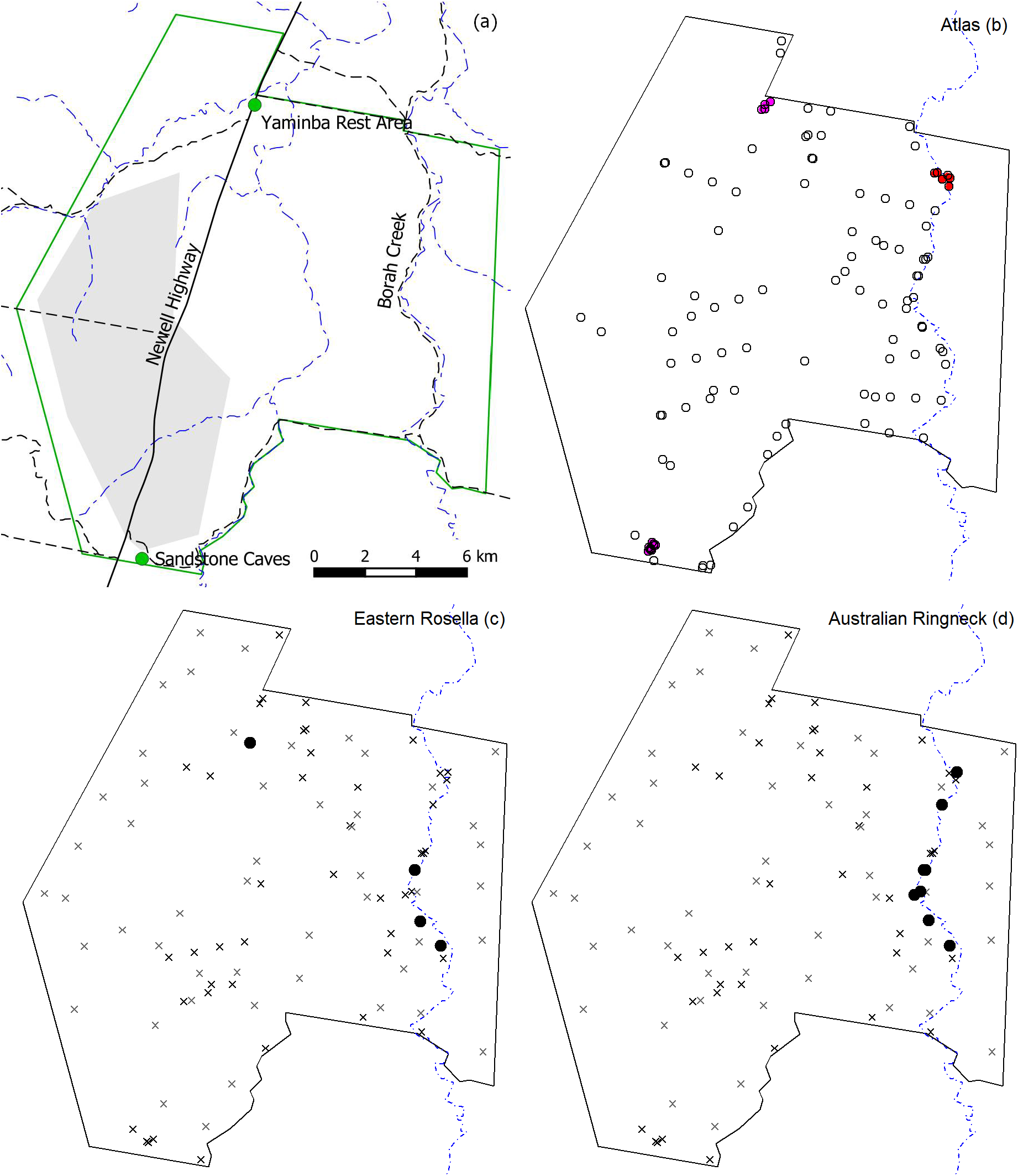
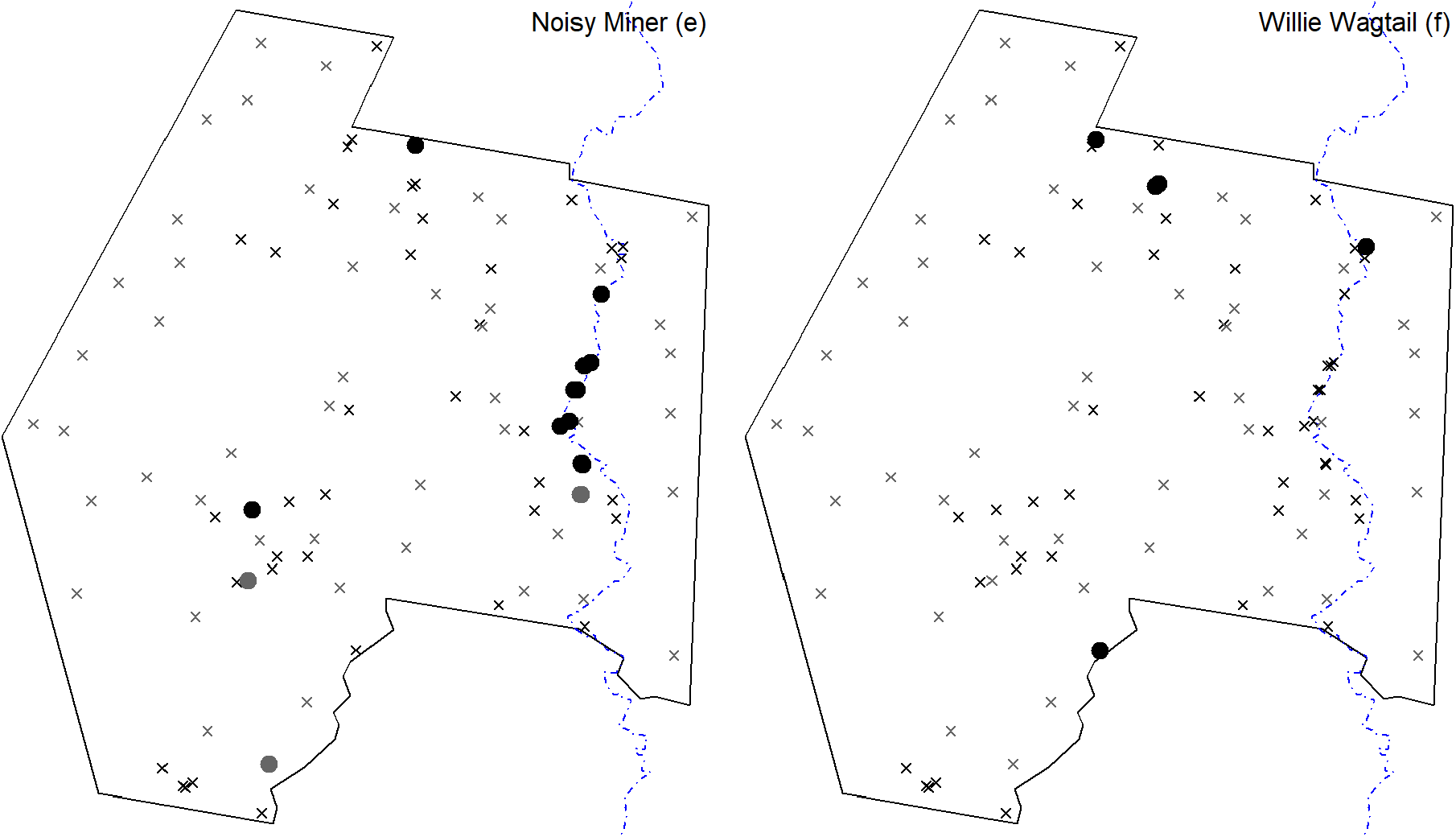
Pilliga study site map (a), atlas spatial sampling pattern (b) (*n* = 108) and species distribution examples (c–f). Solid and dashed black lines in (a) are sealed and unsealed roads respectively (minor tracks are not shown), blue lines are intermittent creek lines and green circles are tourist areas. The grey polygon in (a) indicates an area that had been burnt at some unknown time preceding the survey. Coloured areas in (b) are clusters that were defined using a kernel density and 75% isopleth. Magenta areas correspond to shared sites in the atlas (BirdLife Australia 2021).

### Survey method

The sampling unit was the atlas recommended 20-minute search of a 200 × 100 m (two-hectare) area (Barrett *et al*. 2003) by a single observer. Two-hectare searches were centred on the sampling point coordinates. Points were located and search dimensions were approximately measured using a handheld GPS receiver (Garmin GPSMAP 62s). All bird species inside the two-hectare search area were counted except for those flying over, which were recorded as present/absent.

### Sampling design

Sample sizes for each study site were 50 atlas points, selected at random, and 50 regular points. Spatial sampling was performed using QGIS version 2.16 (QGIS Development Team 2016):

1. A base map of the site was made from screenshots of the atlas website (BirdLife Australia 2017). Distinct survey points can be seem by zooming in. Any repeated sampling of points was unknown and ignored in this study.
2. Between 7–12 control points for georeferencing the base map were obtained from Google Maps (Google 2017). The atlas website plotted points on Google Maps.
3. Control points, in geographic coordinates (latitude and longitude) were projected to Universal Transverse Mercator (UTM). UTM was selected for local accuracy and ease of navigation, with Easting and Northing coordinates in metres.
4. The base map was georeferenced with the UTM control points.
5. Between 9–13 GPS survey points were used to check the accuracy of the georeferenced image. Mean errors were in the range 9–34 m and much smaller than the two-hectare search dimensions.
6. The boundary of the site was digitised using the Google Maps boundary or a polygon was drawn around preexisting atlas points.
7. Atlas points inside the site boundary were digitised. Any repeated sampling of points in the atlas was unknown and not included in this study.
8. A random sample of 50 atlas points were selected, saved as a GPX file and copied to the GPS receiver.
9. A regular sample of 50 points inside the site boundary, with random Easting and Northing offsets, was selected, saved as a GPX file and copied to the GPS receiver. Regular sampling was used to maximise site coverage and minimise spatial autocorrelation.

Surveying occurred from dawn on each day until bird activity was noticed to decline, which usually occurred around mid-morning. Surveying was not performed in windy or rainy conditions, when it is difficult to detect, identify and count birds. Closely-spaced points, less than approximately 400 m apart, were surveyed on different days to reduce temporal autocorrelation in the data.

### Statistical analysis

The analysis examined the three study sites separately. Atlas spatial sampling patterns were quantified using Clark-Evans tests, where *R* = 1 indicates a random pattern, *R* < 1 indicates an aggregated pattern and *R* > 1 indicates a regular pattern (Clark and Evans 1954). These point patterns were bounded by site boundaries and edge corrections were not applied.

Clusters of atlas points were identified using kernel density estimator methods like those used to define monitoring sites for Australian Bird Indices (BirdLife Australia 2015). A kernel density is a bivariate function that gives the probability density that a survey is found at a point according to its spatial coordinates. A cluster is defined as the minimum area for which the sampling probability is equal to some specified value. Kernel densities were computed with an Epanechnikov kernel, 200 m bandwidth and 50 × 50 m cells. Clusters were identified using a 50% or 75% isopleth and only those with at least four survey points were accepted (Ehmke and Herman 2014).

Reporting rates were used to compare atlas sample and regular sample observations. Presence-absence data are closely-linked to spatial sampling patterns and Australian Bird Indices are based on reporting rates (Cunningham and Olsen 2009). Counts are an additional source of variation and counts are optional in the atlas (Barrett *et al*. 2003). Reporting rates were calculated only for diurnal forest birds, *i,e*. excluding any waterbirds, aerial foragers, nocturnal birds and raptors. Overflying birds were also excluded from reporting rate calculations because they were not using the habitat in the two-hectare search area (Szabo *et al*. 2012).

Reporting rate comparisons focused on individual species and atlas sample/regular sample ratios. Ratios greater than at least two-fold (Szabo *et al*. 2012) for ‘non-rare’ species, *i.e*. with at least four records in either survey (reporting rate ≥ 0.08), indicated results that warranted further scrutiny. Four records was the minimum required for statistical significance.

Individual species data can be cast as a two-by-two contingency table (atlas/regular sample × presence/absence). Confidence intervals for reporting rate ratios (‘relative risk’) were computed using the unconditional score statistic method (Agresti and Min 2002). A confidence interval that excludes unity provides evidence that the reporting rates are different.

Spatial autocorrelation, *i.e*. where observations from closely spaced locations are often more similar than are those from widely spaced locations, violates the assumption of independent outcomes for the binomial distribution. Spatial autocorrelation was checked using spline correlograms (Bjørnstad and Falck 2001), which were computed for the combined atlas and regular sample datasets. Statistical comparisons are not reported where the correlogram *y*-intercept (the extrapolated correlation at zero distance) was greater than or equal to 0.2. The theoretical maximum correlation equals one. Spatial statistical methods were investigated and, while they could be useful for reducing bias that results from clustered sampling, this study was concerned with the direct estimation of bias in the atlas sample with reference to the regular sample. Finally, large reporting rate ratios were interpreted with reference to distribution maps and habitat observations.

All statistical analyses were performed using the software R version 4.0.4 (R Core Team 2021). Spatial point pattern statistics were computed using the R package *spatstat* version 2.0-1 (Baddeley *et al*. 2015). Spatial clusters were identified using *adehabitatHR* version 0.4.19 (Calenge 2006). Spline correlograms were computed using *ncf* version 1.2-9 (Bjørnstad 2020). Risk ratio confidence intervals were computed using *PropCIs* version 0.3-0 (Scherer 2018). Figures were prepared using *ggplot2* version 3.3.3 (Wickham 2016).

## RESULTS

### Killawarra

The atlas spatial sampling pattern for Killawarra showed clusters of points at the camp ground and at dams (Fig. 1b) (*n* = 154, Clark-Evans *R* = 0.82, *p* < 0.001). The five clusters were between 13–56 Ha in area, they contained between 5–30 points and 64 points in total (42% of 154).

There were 35 non-rare species (reporting rate ≥ 0.08 in either the atlas sample or the regular sample sample) and 13 of those (37%) had at least two-fold differences in atlas sample/regular sample survey reporting rate ratios (Fig. 4a). Two of those differences were statistically significant (6% of 35) and three were not tested because of spatial autocorrelation (9% of 35) (Table 1).

**Table 1.**
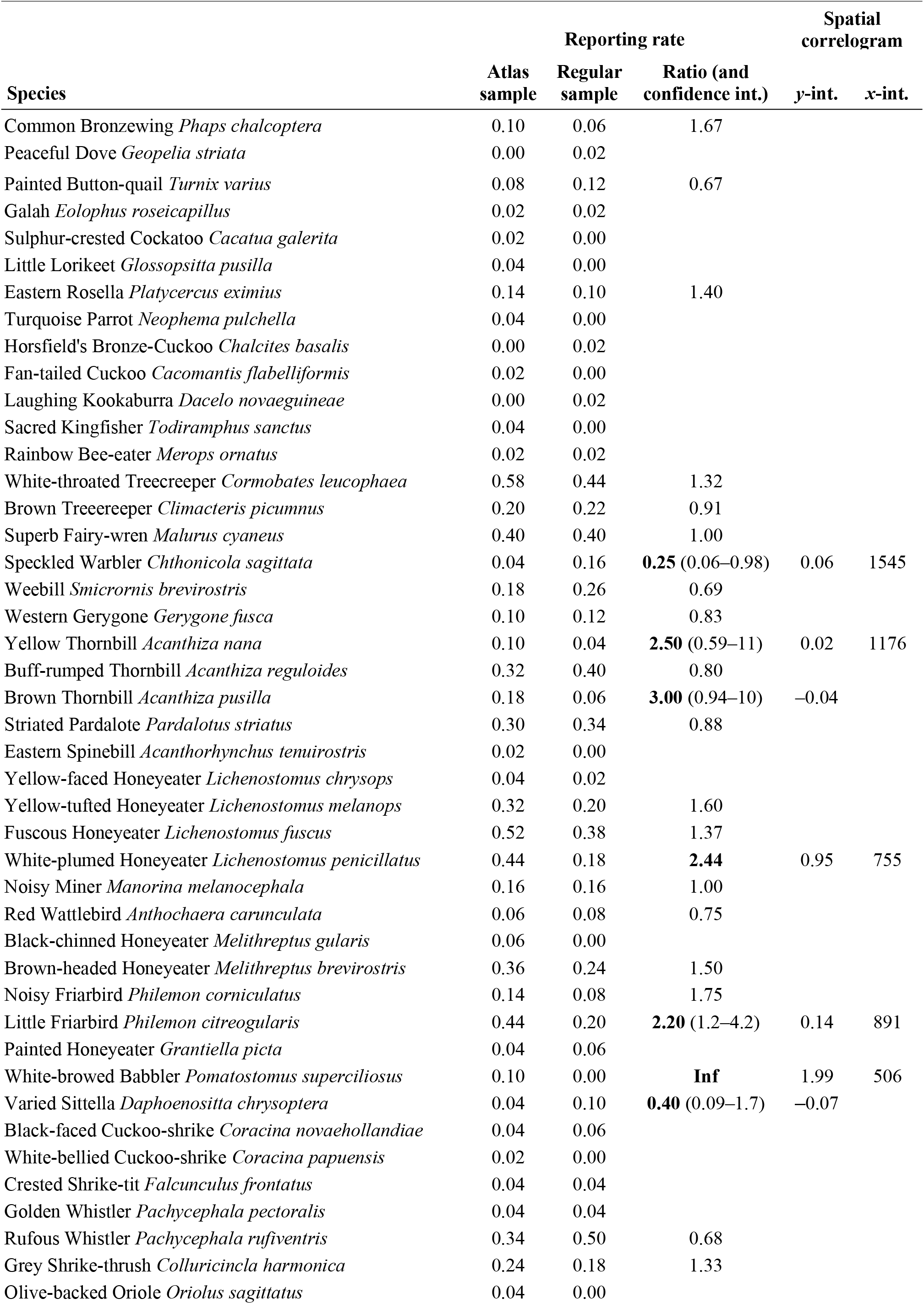
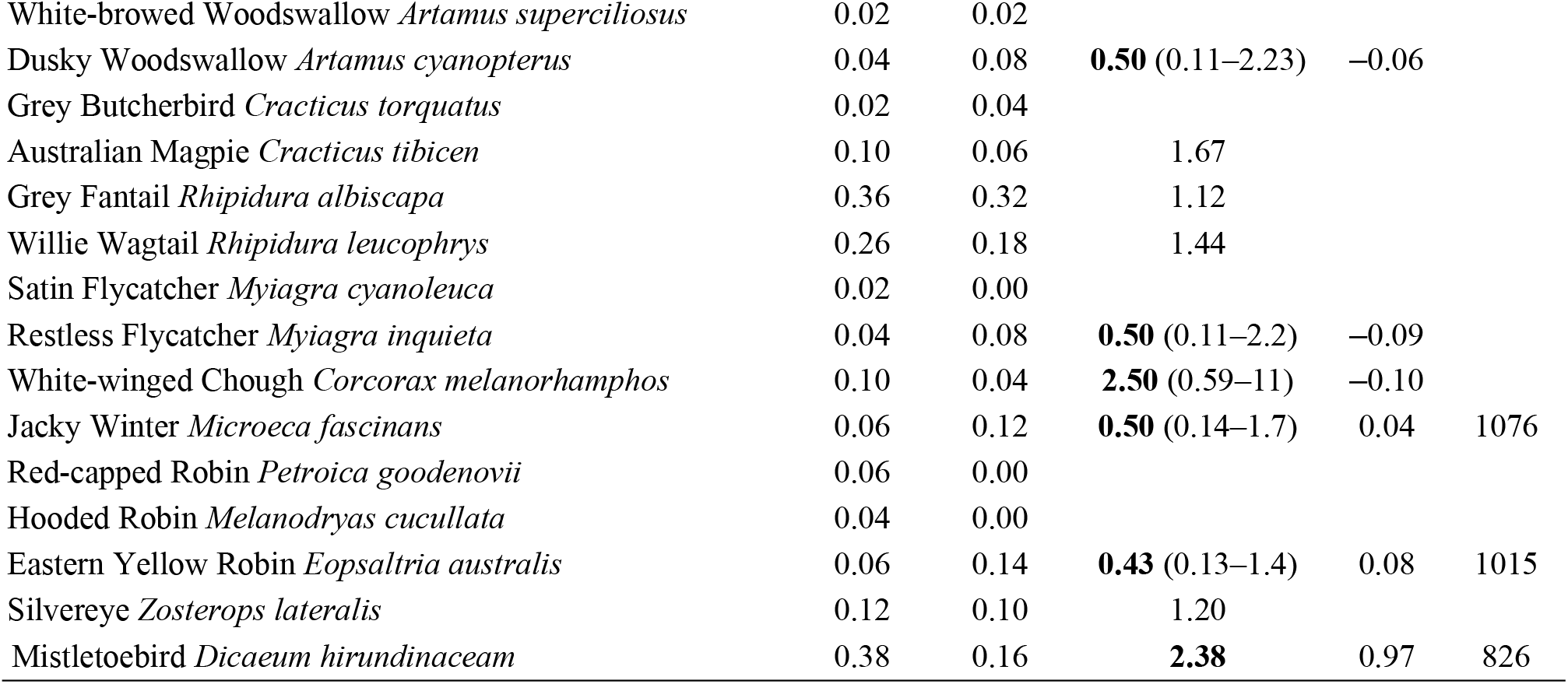
Atlas sample and regular sample reporting rate comparisons for Killawarra. Ratios are not reported for rare species (with both reporting rates < 0.08). At least two-fold differences are in bold. Confidence intervals are not reported for species with spatially correlated observations (correlogram *y-*intercept > 0.2). The correlogram *x*-intercept is the distance at which observations are no more similar than that expected by chance. The correlogram *x*-intercept is not reported if the *y-*intercept is negative.

**Figure 4.**
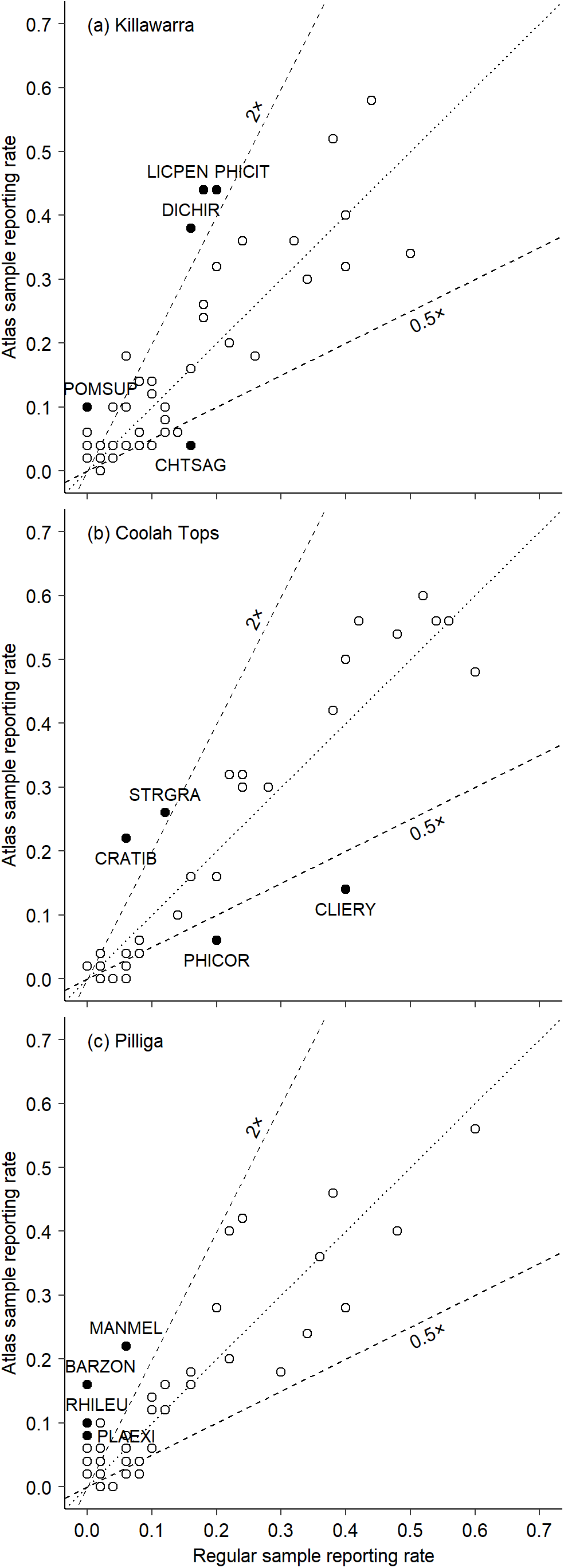
Reporting rate comparison for Killawarra (a), Coolah Tops (b) and Pilliga (c). Dotted lines indicate a 1:1 relationship and dashed lines indicate two-fold differences. Black-filled circles identify species that are noted in the Results. CHTSAG is Speckled Warbler, LICPEN is White-plumed Honeyeater, PHICIT is Little Friarbird, POMSUP is White-browed Babbler and DICHIR is Mistletoebird in (a). CLIERY is Red-browed Treecreeper, PHICOR is Noisy Friarbird, CRATIB is Australian Magpie and STRGRA is Pied Currawong in (b). PLAEXI is Eastern Rosella, BARZON is Australian Ringneck, MANMEL is Noisy Miner and RHILEU is Willie Wagtail in (c). Reporting rates and statistical comparisons are detailed in Tables 1–3.

The atlas sample overestimated reporting rates for species that aggregated at the camp ground and/or dams including White-plumed Honeyeater *Lichenostomus penicillatus*, Little Friarbird *Philemon citreogularis*, White-browed Babbler *Pomatostomus superciliosus* (a family of babblers was resident at the camp ground) and Mistletoebird *Dicaeum hirundinaceam* (there were fruiting mistletoes at the camp ground) (Figs 1c–e). The atlas sample underestimated the reporting rate for Speckled Warbler *Chthonicola sagittata*, which did not frequent those areas (Fig. 1f).

### Coolah Tops

The atlas spatial sampling pattern for Coolah Tops showed clusters of points at camp grounds and other tourist areas in the west, relatively few points in the east and a gap in the middle of the forest (Fig. 2b) (*n* = 116, Clark-Evans *R* = 0.58, *p* < 0.001). The six clusters were between 14–66 Ha in area, they contained between 4–25 points and 60 points in total (52% of 116).

There were 23 non-rare species and five of those (22%) had at least two-fold differences in atlas sample/regular sample reporting rate ratios (Fig. 4b). Two of those differences were statistically significant (9% of 23) and one was not tested because of spatial autocorrelation (4% of 23) (Table 2).

**Table 2.**
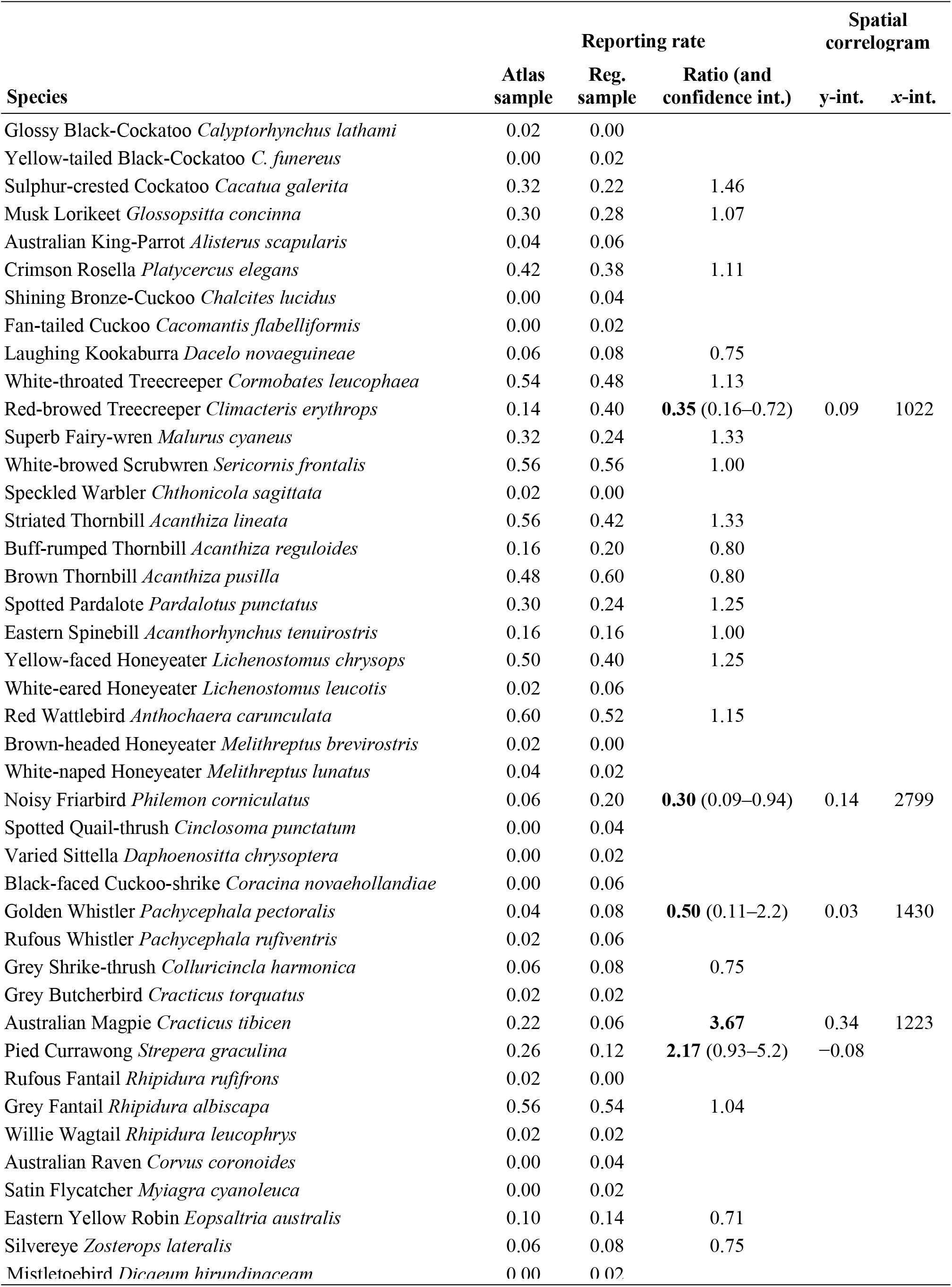
Atlas sample and regular sample reporting rate comparisons for Coolah Tops.

The atlas sample overestimated reporting rates for two species that were associated with more open habitats and/or camp grounds including the Australian Magpie *Cracticus tibicen* and Pied Currawong *Strepera graculina* (Figs 2c–d). The atlas sample underestimated reporting rates for Red-browed Treecreeper *Climacteris erythrops*, which was apparently associated with stringybark forest along ridges, and Noisy Friarbird *Philemon corniculatus* (Figs 2e–f).

### Pilliga

The atlas spatial sampling pattern for the Pilliga study site looked well-dispersed, including lines of regularly spaced points (Fig. 3a). Nonetheless, there were clusters of points at tourist areas and linear patterns along Borah Creek (*n* = 108, Clark-Evans *R* = 0.71, *p* < 0.001). The three clusters were between 20–32 Ha in area, they contained between 4–9 points and 19 points in total (18% of 108).

There were 26 non-rare species and seven of those (26%) had at least two-fold differences in atlas sample/regular sample reporting rate ratios (Fig. 4c). One of those differences was statistically significant (4% of 26) and three were not tested because of spatial autocorrelation (12% of 26) (Table 3).

**Table 3.**
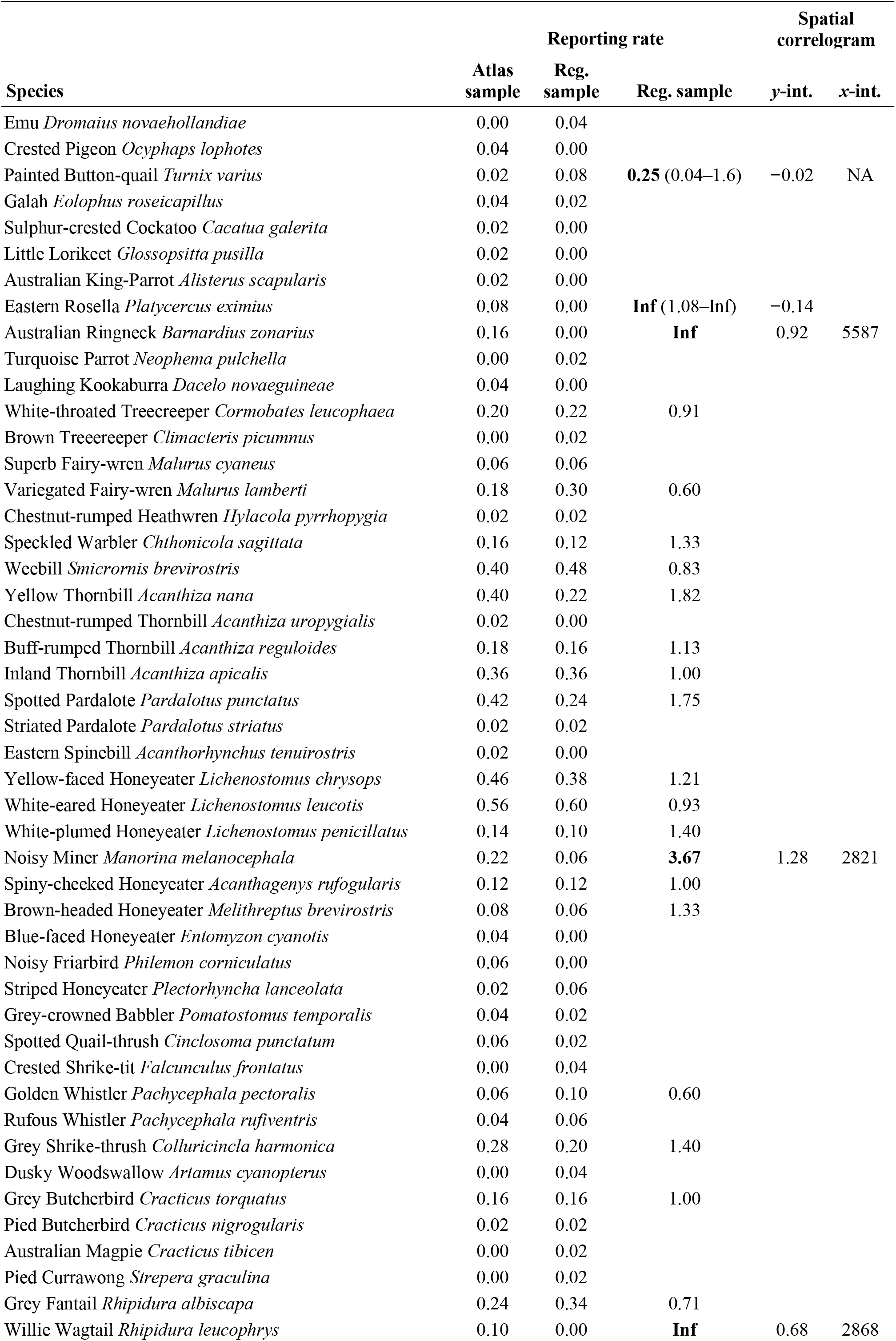
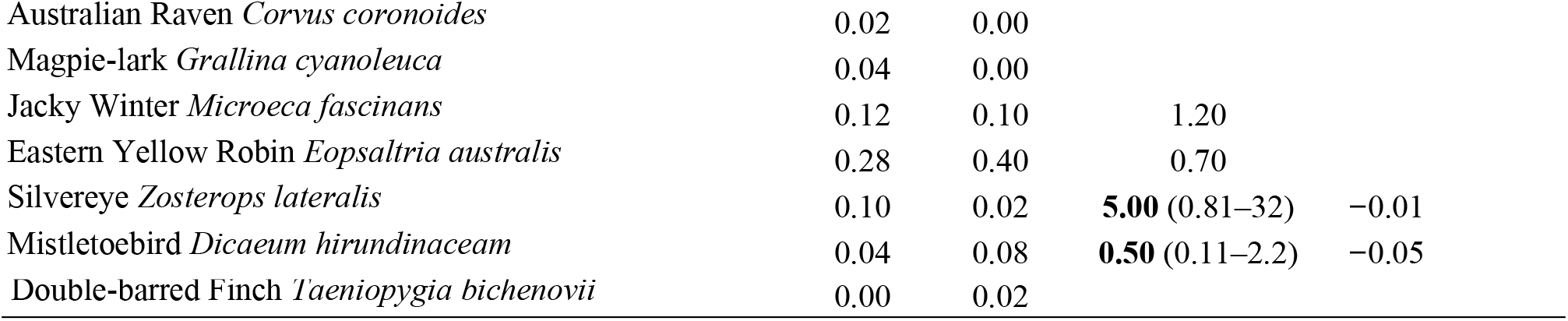
Atlas sample and regular sample reporting rate comparisons for the Pilliga study site.

The atlas sample overestimated reporting rates for species that were associated with the grassy woodland habitat along Borah Creek including Eastern Rosella *Platycercus eximius*, Australian Ringneck *Barnardius zonarius*, Noisy Miner *Manorina melanocephala* (Figs 3c–e) and the rare Crested Pigeon *Ocyphaps lophotes* and Blue-faced Honeyeater *Entomyzon cyanotis*.

## DISCUSSION

Convenience and subjective sampling was evident in atlas spatial sampling patterns for all three sites in this study. Regular and random sampling designs could be criticised for missing special habitat features, *e.g*. creek lines and dams, however patch-scale mean abundance estimates were the objective of this study. If required, complex survey designs can incorporate special habitat features, *e.g*. via stratified sampling or cluster sampling, and, because every sampling unit has a known, non-zero probability of selection, result in unbiased mean and variance estimates (Lemeshow and Levy 1999).

At least two-fold differences in atlas/regular sample reporting rates were detected for between 13–15% of non-rare bird species. These numbers are higher than expected by chance alone at the five percent significance level. More species with large reporting rate differences, especially rare birds, could have been detected with larger sample sizes, *e.g*. 28% of species in the Mount Lofty Ranges evaluation (Szabo *et al*. 2012).

Sampling biases occur at all spatial scales in the atlas (Barrett *et al*. 2003; BirdLife Australia 2015). Spatial sampling bias has been found to strongly affect reporting rates for a substantial proportion of woodland and forest bird species at both local (this study) and regional scales (Szabo *et al*. 2012), the identity of those species cannot be predicted from habitat associations, *i.e*. they were not all edge and open habitat species (*contra* Szabo *et al*. 2012), and large sample sizes increase precision but do not reduce bias (*e.g*. Szabo *et al*. 2012).

Clusters in atlas spatial sampling patterns at Killawarra and Pilliga corresponded with a number of ‘shared sites’ in the atlas that are assumed to be monitoring sites for Australian Bird Indices. These shared sites were located at a tourist areas and dams and were not representative of those forests. Also, between 48–72% of atlas points were not located within any clusters and so may not be useful for Australian Bird Indices. Similarly, the Mount Lofty Ranges evaluation started with 3700 atlas surveys and selected 554 (15%) that were in eucalypt woodland in spring–summer (Szabo *et al*. 2012). Volunteers would be disappointed to learn that so many surveys are being filtered out.

The atlas would benefit from a sampling design and, because the Australian continent is vast and the bird watcher population size is small, purposive selection of monitoring sites would be more efficient than random or other probability sampling designs. The coordinates of these monitoring sites could be provided as GPX files that can be uploaded to GPS navigation devices and used to guide volunteers to recommended sites.

## ACKNOWLEDGEMENTS

None at present.

